# Structural connectivity predicts individual perceived stress

**DOI:** 10.1101/2023.12.07.570637

**Authors:** Toluwani Joan Amos, Olusola Bamisile, Zhenlan Jin, JunJun Zhang, Li Ling

## Abstract

Many previous studies have investigated the neural mechanisms of perceived stress using either task or resting-state functional connectomes. However, to date, the structural connectivity predictors of individual perceived stress remain unknown. In this study, using connectome-based predictive modeling with a leave-one-out cross-validation framework in a sample of 100 unrelated healthy young adults, we show that individual differences in perceived stress can be reliably predicted from their structural connectivity. The obtained results show that perceived stress could be predicted from the interaction of visual association, and motor and sub-cortical networks. This present work highlights that structural connectivity can be used to investigate the neural mechanism of PS in healthy populations.

## I. Introduction

Every healthy individual experiences stress in different variations and the subjective measure of individual stress levels can be quantified using perceived stress (PS), which is the rate at which life’s situations are estimated as unpredictable and uncontrollable(S. Wang et al., 2019). Increased PS levels have been associated with illnesses such as cardiovascular diseases, and stroke (Levine, 2022), as well as psychiatric disorders such as anxiety and depression (Oathes et al., 2015).

Previous functional studies that have investigated the neural mechanisms of PS have highlighted the involvement of the limbic systems and prefrontal cortex (Hermans et al., 2014; van Oort et al., 2017). Others have also implicated limbic regions such as the hippocampus and parahippocampus to play vital roles in explaining the processes involved in stress (Albert et al., 2015; Chang & Yu, 2019). Generally, these functional network studies have demonstrated that stress response can be linked with alterations in multiple brain networks.

Previous research has significantly contributed to promoting the general understanding of the neural basis of stress.

However, no study has investigated the structural connectivity predictors involved in predicting PS at the individual level in healthy individuals. This research aims to address this research gap using individual structural connectomes and a cross-validated framework known as connectome-based predictive modeling (Shen et al., 2017).

Connectome-based predictive modeling (CPM) is a novel machine learning method developed with built-in cross-validation for extracting and summarizing the most relevant features from the brain connectivity data, which is used to construct the prediction models. The ability to predict individual behavior from whole-brain functional connectivity is an important development in research, as it shifts the focus from group analyses to single-subject analysis by emphasizing inter-subject differences. CPM has been successfully used to predict various emotion-related behavioral scores, such as individual trait anxiety (Z. Wang et al., 2021), loneliness (Feng et al., 2019), and self-reported emotional intelligence scores (He et al., 2023).

In this study, we implemented a CPM to test whether a whole-brain connectome based on structural connectivity can reliably predict PS in novel participants within a sample of young healthy adults.

## II. Methods

### A. Dataset description

T1-weighted and diffusion MRI images from 100 unrelated individuals from the Human Connectome Project-Young Adults (Van Essen et al., 2013). T1 anatomical images were acquired using the 3D magnetization-prepared rapid gradient echo sequence (MPRAGE) with 0.7 x 0.7 x 0.7 mm^3^ voxel size, TR/TE = 2400/2.14 ms, and flip angle = 8 degrees. The dMRI data was acquired on a customized Siemens Magnetom Skyra 3T MRI system using a multi-band pulse sequence. The protocol for the diffusion imaging comprised 3 diffusion-weighted shells (b=1000, 2000, and 3000 s/mm2). To ensure distortion correction, all images were additionally acquired with reversed phase encoding. Other diffusion-weighted imaging parameters which were used include 145 x 145 matrix, 174 slides, 1.25 x 1.25 x 1.25 mm3 voxel size, TR/TE = 5520/89.5ms.

### B. Perceived stress measurements

Perceived stress (PS) is defined by the individual perceptions about the nature of events and their relationship to the values and coping resources. The individual PS was quantified using the National Institute of Health toolbox perceived stress survey. The PS survey is a self-reported measure that assesses how unpredictable, uncontrollable, and overloaded individuals find their lives (Babakhanyan et al., 2018). Previous research has highlighted that the PS scale has high reliability and is a sensitive measure of subjective stress levels across diverse samples (Ng, 2013).

### C. Structural connectome construction

First, further pre-processing was performed on the HCP data using the MRtrix3 software package (http://www.mrtrix.org) on a flexible, light-weight, scalable, and out-of-the-box analysis environment called NeuroDesk (Renton et al., 2023). The pre-processing steps that we applied to the HCP data included, bias-field correction, followed by multi-shell multi-tissue constrained spherical deconvolution (CSD) (Wilson et al., 2021) to model the white matter, grey matter, and the cerebrospinal fluid using a maximum harmonic degree Lmax=8. Second, this initial pre-processing was followed by the generation of the structural connectome for each subject, which is composed of three major steps such as tractogram construction, spherical-deconvolution informed filtering of tractograms (SIFT2), and connectome generation. Before the tractogram construction, the T1w anatomical image was segmented into five tissue types such as grey matter, sub-cortical grey matter, white matter, cerebrospinal fluid, and, pathological tissue. Then, the segmented anatomical image was co-registered with the diffusion image using FMRIB software library (FSL) 5.0 and Analysis of functional neuroimages (AFNI) 21.2.0 software implemented in the NeuroDesk environment. Next, a seed boundary that separates the grey from the white matter (grey matter/white matter interface) was generated which serves as the seeds for the streamlines. In the tractogram construction, 10 million probabilistic streamlines using the 2nd order Integration over Fibre Orientation Distributions algorithm (Gutierrez et al., 2020) and anatomically-constrained tractography (ACT), FOD amplitude threshold 0.06, step size which is half of the voxel size, length of 5-300mm and backtracking was performed. Backtracking helps to check for truncated tracks and ensures re-tracking if a poor structural termination is encountered (Tuladhar et al., 2020). Next, we filtered the tracks using the Spherical-deconvolution Informed Filtering of Tractograms (SIFT2) algorithm, which aids in reducing the overall track count and yields more biologically meaningful estimates of structural connection density. Finally, we built the whole brain structural connectome matrix using the generated white matter tracks and the parcellated nodes. The nodes of the connectome matrix are the parcellated ROIs, and the values of the matrix are the white matter connectivity strengths among the ROIs. From each subject’s tractogram, an individual weighted structural connectome, which in principle, provides a more powerful characterization of biological properties was computed using the Shen atlas. The structural connectome comprised 268 regions of interest (cortical and sub-cortical regions) and the connection strengths were calculated by summing the weights of the relevant streamlines.

### D. Connectome-based predictive modeling (CPM)

We employed connectome-based predictive modeling with a leave-one-out cross-validation approach in 100 unrelated subjects to determine whether whole-brain structural connectivity could predict individual perceived stress in the healthy adult cohort. For feature extraction, a vector of behavioral responses for each participant was Spearman correlated with the 35,778 edges in the functional connectivity matrix. Next, we retained the edges that were significantly positively and negatively correlated with the behavioral score (p<0.01). The weights of the positive and negative edges were summed to obtain a positive and negative network strength, respectively. A combined network strength was also constructed from the difference in the positive and negative network strength. Then, we used linear regression for the model building to fit the single-subject summary network strength values from the positive, negative, and combined networks with the PS scores. In the prediction stage, the obtained regression coefficients from the trained models were used to predict the target variable for the left-out participant. The entire analysis was repeated 100 times until each participant had served as the test participant. The Spearman’s rank correlation between the observed and predicted PS scores was used in evaluating the model’s predictive power.

### E. Anatomical distribution of predictive edges

We defined whole-brain network masks to identify the significant predictive functional edges during the feature extraction step of CPM. These masks contained the edges that appeared in every iteration of the leave-one-out cross-validation. Thus, the models derived from the structural connectivity yielded two separate masks, which included the edges that showed significant positive and negative correlations with the PS scores, respectively. Next, to visualize the significant functional edges in each respective mask, which was arranged according to the 268 nodes from the Shen brain atlas, we utilized the interactive Bioimage Suite tool https://www.nitrc.org/projects/bioimagesuite/.

## III. Results

### A. Predicting individualized PS scores from whole-brain structural connectivity

The whole brain structural connectivity succeeded in predicting the individual PS scores using CPM and the leave-one-out-cross-validation framework. Specifically, in predicting the PS scores, only the negative network model (*r* = .306, *p* = .002) and combined network model (*r* = .358, *p* <0.001) yielded significance, as shown in Fig. 1D.

**Fig. 1.**
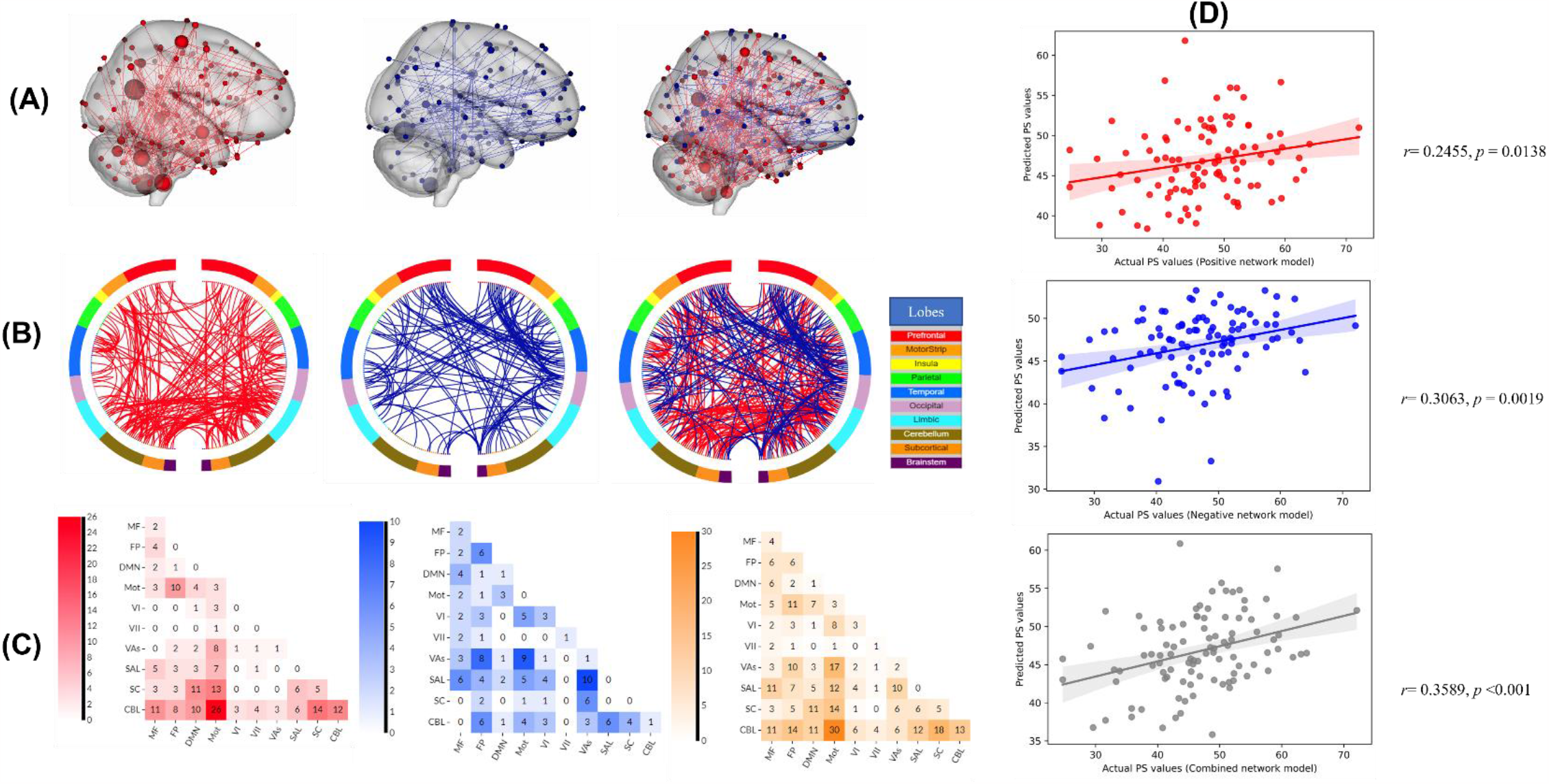
Structural connectivity predicts individual perceived stress scores. (A) Glass brain plots showing the right view of the significant functional edges across the predictive models (B) Circle plots showing the significant connections from the brain lobes contributing to the predictive models. Positive network: ted (left). Negative network: blue (middle), stun of positive and negative edges: orange (right). (C) Matrix plot indicating the highest contributing networks. (D). Spearman correlation plot between the actual and predicted ΔMRT for the positive, negative, and combined network models respectively.

### B. Network neuroanatomy in the prediction of individual PS scores

Next, we investigated the top contributing networks in the predictive models. We observed that the top contributing networks in predicting the individual PS scores were mainly from the visual association-motor, salience-visual association network interactions in the positive network and network interactions between cerebellum-motor and cerebellum-subcortical areas (Fig. 1A-C).

## IV. Discussion

In this project, we set out to test for the first time, if it was possible to predict the individual perceived stress scores from their structural connectivity. Using the CPM framework, we were able to build predictive models to achieve this aim. Also, we observed distinct network contributions in predicting the individual PS scores from the positive and combined network models. A substantial contribution of the motor network interacting with other brain networks, such as the visual network, cerebellum, and sub-cortical areas was seen. A recent study that used resting state functional networks in predicting perceived stress(Liu et al., 2021) highlighted the major role of the default mode network interacting with other brain networks in stress processes (Chang & Yu, 2019; Maron-Katz et al., 2016). However, from our study using structural connectivity, the predictors were mainly from interactions between salience-visual association, cerebellum-motor, and cerebellum-subcortical network areas. This suggests that the neural contributions in explaining perceived stress might be different from the functional and structural connectome perspectives and further investigation is needed to clarify this narrative.

## V. Conclusion

In sum, this study demonstrates that individual PS scores in a sample of healthy young adults can be reliably predicted from their structural whole-brain functional connectivity.

